# Identifying biomarkers of anti-cancer drug synergy using multi-task learning

**DOI:** 10.1101/243568

**Authors:** Nanne Aben, Julian R. de Ruiter, Evert Bosdriesz, Yongsoo Kim, Gergana Bounova, Daniel J. Vis, Lodewyk F.A. Wessels, Magali Michaut

**Affiliations:** Division of Molecular Carcinogenesis, Oncode Institute, Netherlands Cancer Institute, Amsterdam 1066CX, The Netherlands; Faculty of EEMCS, Delft University of Technology, Delft 2628CD, The Netherlands; Division of Molecular Pathology, Oncode Institute, Netherlands Cancer Institute, Amsterdam 1066CX, The Netherlands; Cancer Genomics Netherlands, Utrecht 3584CT, The Netherland; Division of Oncogenomics, Oncode Institute, Netherlands Cancer Institute, Amsterdam 1066CX, The Netherlands

## Abstract

Combining anti-cancer drugs has the potential to increase treatment efficacy. Because patient responses to drug combinations are highly variable, predictive biomarkers of synergy are required to identify which patients are likely to benefit from a drug combination. To aid biomarker identification, the DREAM challenge consortium has recently released data from a screen containing 85 cell lines and 167 drug combinations. The main challenge of these data is the low sample size: per drug combination, a median of 14 cell lines have been screened. We found that widely used methods in single drug response prediction, such as Elastic Net regression per drug, are not predictive in this setting. Instead, we propose to use multi-task learning: training a single model simultaneously on all drug combinations, which we show results in increased predictive performance. In contrast to other multi-task learning approaches, our approach allows for the identification of biomarkers, by using a modified random forest variable importance score, which we illustrate using artificial data and the DREAM challenge data. Notably, we find that mutations in MYO15A are associated with synergy between ALK / IGFR dual inhibitors and PI3K pathway inhibitors in triple-negative breast cancer.

**Author summary:** Combining drugs is a promising strategy for cancer treatment. However, it is often not known which patients will benefit from a particular drug combination. To identify patients that are likely to benefit, we need to identify biomarkers, such as mutations in the tumor’s DNA, that are associated with favorable response to the drug combination. In this work, we identified such biomarkers using the drug combination data released by the DREAM challenge consortium, which contain 85 tumor cell lines and 167 drug combinations. The main challenge of these data is the extremely low sample size: a median of 14 cell lines have been screened per drug combination. We found that traditional methods to identify biomarkers for monotherapy response, which analyze each drug separately, are not suitable in this low sample size setting. Instead, we used a technique called multi-task learning to jointly analyze all drug combinations in a single statistical model. In contrast to existing multi-task learning algorithms, which are black-box methods, our method allows for the identification of biomarkers. Notably, we find that, in a subset of breast cancer cell lines, *MYO15A* mutations associate with response to the combination of ALK / IGFR dual inhibitors and PI3K pathway inhibitors.

## Introduction

Combining drugs is a promising strategy for cancer treatment, as drug combinations can increase the efficacy of treatment. For example, Prahallad *et al.* (2012) [17] have shown that combining a BRAF inhibitor with an EGFR inhibitor shows synergy in *BRAF* mutant colorectal cancer. However, for most drug combinations it is not known what subset of patients will respond. By identifying biomarkers (e.g. mutations in the tumor’s DNA that are associated with a favorable response to the drug combination), the selection of a given patient’s treatment can be improved. To facilitate biomarker identification, data from a large-scale drug combinations screen were recently released as part of the AstraZeneca-Sanger DREAM challenge [14], containing 85 cell lines with their response to 167 drug combinations.

While the data from this screen can provide information on potential biomarkers of synergy, it is not yet clear what is the best way to identify them. In the context of single drug response prediction, the default approach is to fit ‘individual models’ that are trained separately per drug. We applied a similar approach here in the context of drug combinations, training ‘individual models’ for each drug combination separately (Fig 1A). However, we show that such an approach is unsuitable for the dataset at hand due to the extremely low sample size: a median number of 14 cell lines have been screened per drug combination.

**Fig 1.**
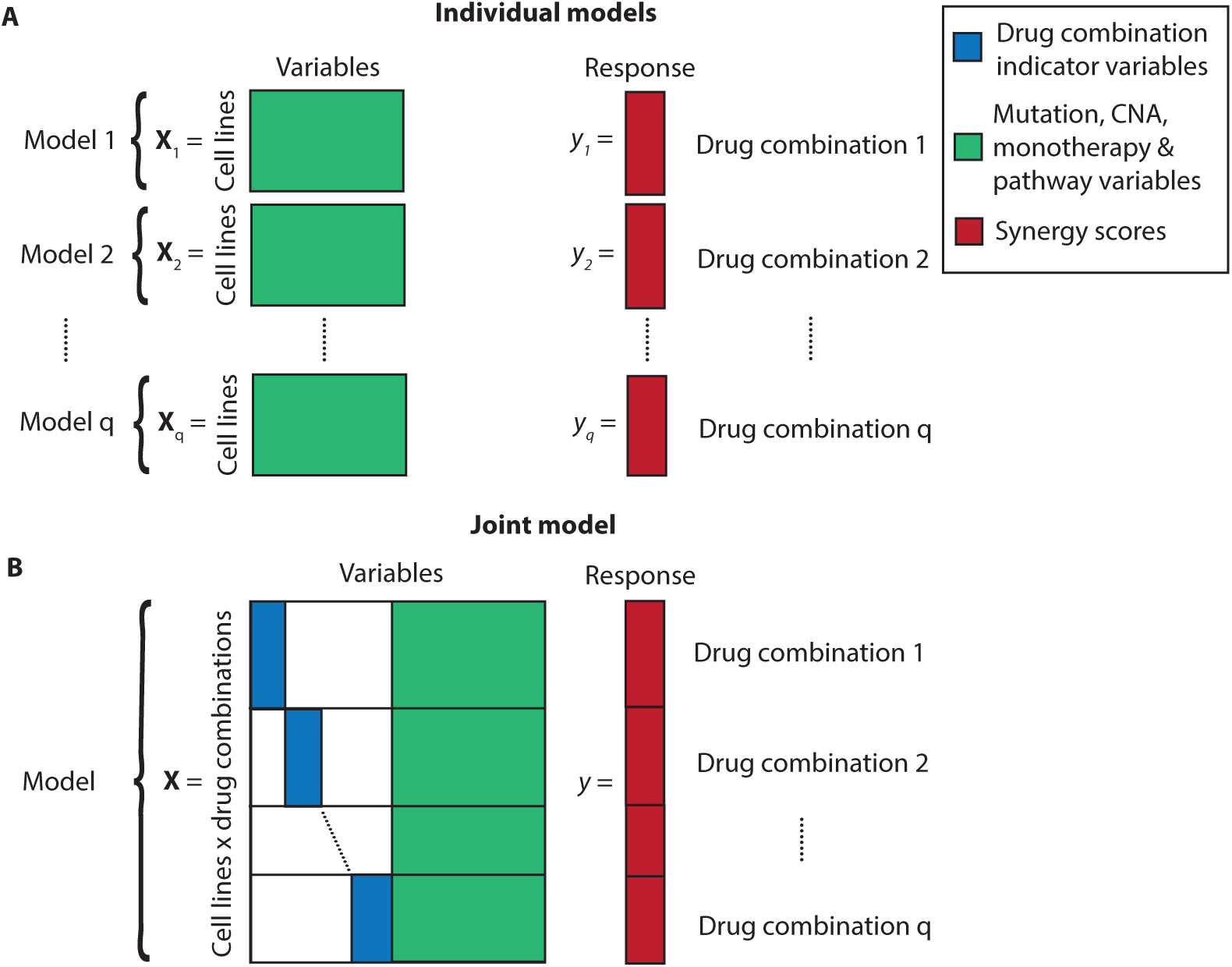
Overview of the models used in this work. A: Graphical representation of *q* individual models, in which a different model is trained independently for each of the *q* drug combinations. B: Graphical representation of the joint model, in which a single model is jointly trained on all *q* drug combinations simultaneously.

We propose to alleviate the problem of low sample size by training ‘joint models’ that use information from all drug combinations simultaneously (Fig 1B). In the literature, this is known as multi-task learning [4,15]. This approach has been employed before in single drug response prediction by Gönen *et al.* (2014) [9], Menden *et al.* (2013) [13] and Yuan *et al.* (2016) [19], and in synergy prediction by other participants in the AstraZeneca-Sanger DREAM challenge [14]. What distinguishes our approach from other multi-task learning approaches is that we are able to identify biomarkers, whereas others have proposed black-box models. Specifically, the Joint Random Forest model we propose is simultaneously trained on all drug combinations, after which we apply our Drug-combination-specific Variable Importance (DVI) score to the trained Joint Random Forest to identify biomarkers of synergy. We provide a Python implementation on our Github (https://github.com/NKI-CCB/multitask_vi/).

We show that the joint model outperforms individual models in terms of predictive performance. Using the joint model together with the DVI, we are able to identify biomarkers of response on both simulated and real data. Finally, we found that *MYO15A* mutations associate with synergy between an ALK / IGFR dual inhibitor and PI3K pathway inhibitors in triple-negative breast cancer.

## Results

### The AstraZeneca-Sanger DREAM challenge data

In order to predict synergy from molecular data, we have used the data from the AstraZeneca-Sanger DREAM challenge (from here on referred to as: DREAM data) [14]. The goal of this community challenge was to create models that predict whether a given drug combination will show synergy in certain cell lines. The DREAM data include 85 cell lines and 167 drug combinations, with a median of 14 cell lines screened per drug combination. The dataset consists of three parts: synergy scores, monotherapy response data and molecular data of the cell lines (e.g. mutations and copy number alteration data).

As the response variable for our model, we used the synergy scores as provided in the DREAM data, which were based on a Loewe additivity model [5,8]. For each cell line, drug combination pair, monotherapy data were available, quantifying the response of a cell line to each individual drug in the drug combination by the 50% Inhibitory Concentration (IC50) or the Area Under the dose-response Curve (AUC). For each cell line, molecular data were provided in the form of mutation, copy number alteration (CNA), methylation and gene expression data. Because of the high dimensionality and the low sample size, we restricted mutations and CNAs to a reduced set of potential driver genes. Finally, we defined ‘pathway rules’ that integrate the mutation and CNA data with information from KEGG [11,12]. More information on how these data were processed is provided in the S1 Text.

We used the monotherapy and the molecular data of the cell lines to predict drug synergy (Fig 1). More formally, we defined the input matrix **X** using 382 mutation, 76 copy number, 23 monotherapy, and 16 pathway rule variables. The response vector *y* was defined using the synergy scores. Each of the input data types explain a part of the synergy and are therefore useful to include in a predictive model. For biomarker identification we focused on genomic variables only, as monotherapy data are unlikely to be useful as clinical biomarkers (this information is typically not available for most drugs for a given patient).

### Per-combination individual models perform poorly

For our initial approach, we used the DREAM data to create ‘individual models’ that are trained separately per drug combination (Fig 1A) (Methods). To test the variability across different prediction methods, the individual models were trained using either Elastic Net, SVM (with RBF kernels) or Random Forest. For each method, predictive performance was assessed using cross-validation with the ‘primary score’ (a weighted average of the correlation between the observed and predicted synergy scores) defined in the DREAM challenge [14] as endpoint.

Overall, the predictive performance of the individual models was low for all methods (0.04 on average) (Fig 2A), most likely due to the extremely low sample size (median of 14 cell lines per combination). We also observed that the predictions from the individual SVM models resulted in negative correlations between the observed and predicted synergy scores (Fig 2A). This is due to a cross-validation artifact that leads to negative correlations when the model is unable to detect structure in the data (S2 Text), which likely mostly affected the SVM due to the high complexity of the RBF kernels.

**Fig 2.**
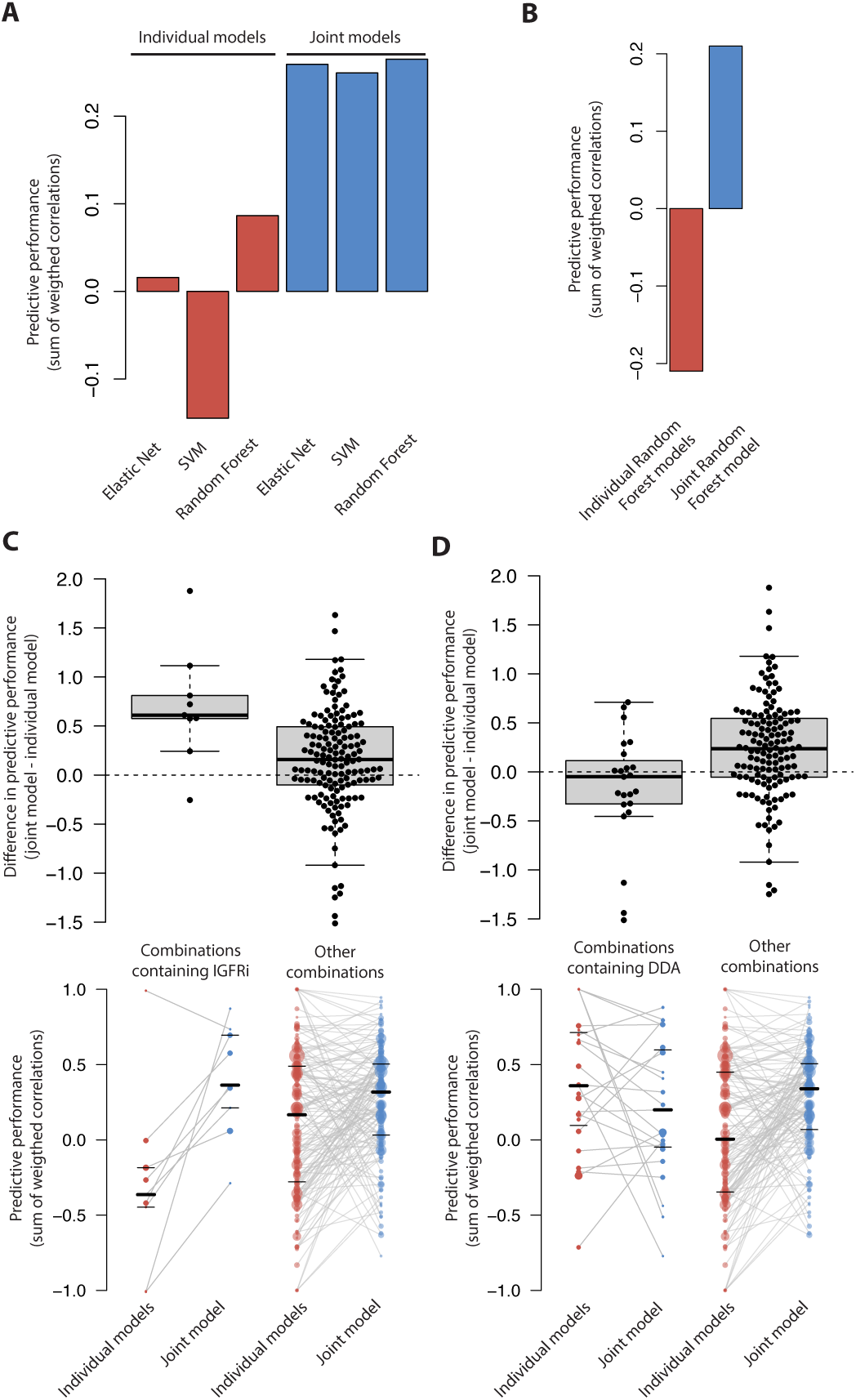
Predictive performance of the joint and individual models. Performance was measured by taking the weighted sum of correlations between the observed and predicted synergy scores. A: Predictive performance stratified by method (Elastic Net, SVM, Random Forest) and model (individual, joint). B: Predictive performance for individual and Joint Random Forest models, assessed using leave-one-cell-line-out cross-validation. C&D: Association between a specific drug class and the difference in predictive performance between individual and joint models (top panel). How the predictive performance changed between the individual and joint models is illustrated in the bottom panel. Each dot represents the predictions for a given drug combination. Predictions for the same drug combination are connected between the individual and the joint models. The size of the dot is proportional to the number of cell lines the model was trained on. From left to right: combinations containing IGFR inhibitors (IGFRi), all other drug combinations (not containing IGFRi), combinations containing DNA damaging agents (DDA), and all other combinations (not containing DDA). Note that the extreme correlations (i.e. correlations close to 1 or -1) can be attributed to small sample size (indicated here by the size of the dot in the bottom panel).

### Simultaneously learning across drug combinations improves predictive performance

To alleviate the low sample size problem, we created ‘joint models’, which are trained on all drug combinations simultaneously, thereby leveraging the information from the entire dataset. Using cross-validation, we found that the joint models achieve higher predictive performance compared to individual models (Fig 2A), regardless of the underlying method (Elastic Net, SVM, Random Forest).

A drawback of the standard cross-validation scheme is that the same cell line can be in different cross-validation folds (but for different drug combinations), which could bias the predictive performance. To test for this, we also performed leave-one-cell-line-out cross-validation, in which all data associated with a given cell line were left out from the training step of a given fold. Overall, we found that joint models were more predictive than individual ones using leave-one-cell-line-out cross-validation too (Fig 2B), ruling out this bias. We also observed that the individual Random Forest models resulted in negative predictive performance in this setting (Fig 2B), whereas the predictive performance was positive using regular cross-validation (Fig 2A). This too can be attributed to the aforementioned cross-validation artifact (S2 Text).

To determine whether the joint models were predictive for specific classes of drug, we grouped the 119 drugs into 19 drug classes and checked whether the difference in predictive performance between the individual or joint models was associated with any of the drug classes. This showed that drug combinations containing IGFR inhibitors are significantly better predicted using the joint model (Mann-Whitney U test, FDR-corrected *p* = 0.047) (Fig 2C). Furthermore, for drug combinations containing DNA damaging agents (DDA), the joint model showed on average no increase in predictive performance (Fig 2D, bottom panel). Compared to the overall increase in predictive performance between individual and joint models, this effect was significant (Mann-Whitney U test, FDR-corrected *p* = 0.036) (Figure 2D, top panel).

To further characterize the joint model predictive performance improvement, we used a simulated dataset and assessed under which conditions joint models outperform individual models using Random Forests. In this simulation, we created a data set of similar size as the DREAM data and then varied the sample size or the number of features (Methods). We found that simultaneously learning across drug combinations was most beneficial in highly underdetermined cases, i.e. when the sample size was low or the number of variables was high (Fig S1). Interestingly, when the number of samples was sufficiently high (e.g. *n* = 100), the individual and joint Random Forest models achieved virtually identical predictive performance.

Altogether, our results show that, for most combinations, joint models obtain a higher predictive performance compared to individual models by simultaneously learning across drug combinations. As the joint Random Forest model obtained the highest predictive performance, we decided to further use this joint model for biomarker identification.

### Joint model Variable Importance scores are not sufficient to identify biomarkers of synergy

In an initial attempt to identify biomarkers of synergy using the joint model, we first computed the Random Forest’s variable importance score (VI), referred to as the joint model VI score (JVI) (Fig 3A). Ranking the variables by their JVI, we identified variables that had a large impact on the prediction of many different drug combinations. We found that the monotherapy variables were the most important variables overall (highest JVI scores in Fig 4) (one-tailed Mann-Whitney U test, *p* = 2.474*e*-16) (S3 Text), followed by pathway rules (one-tailed Mann-Whitney U test, *p* = 6.248*e*-06) (S4 Text).

**Fig 3.**
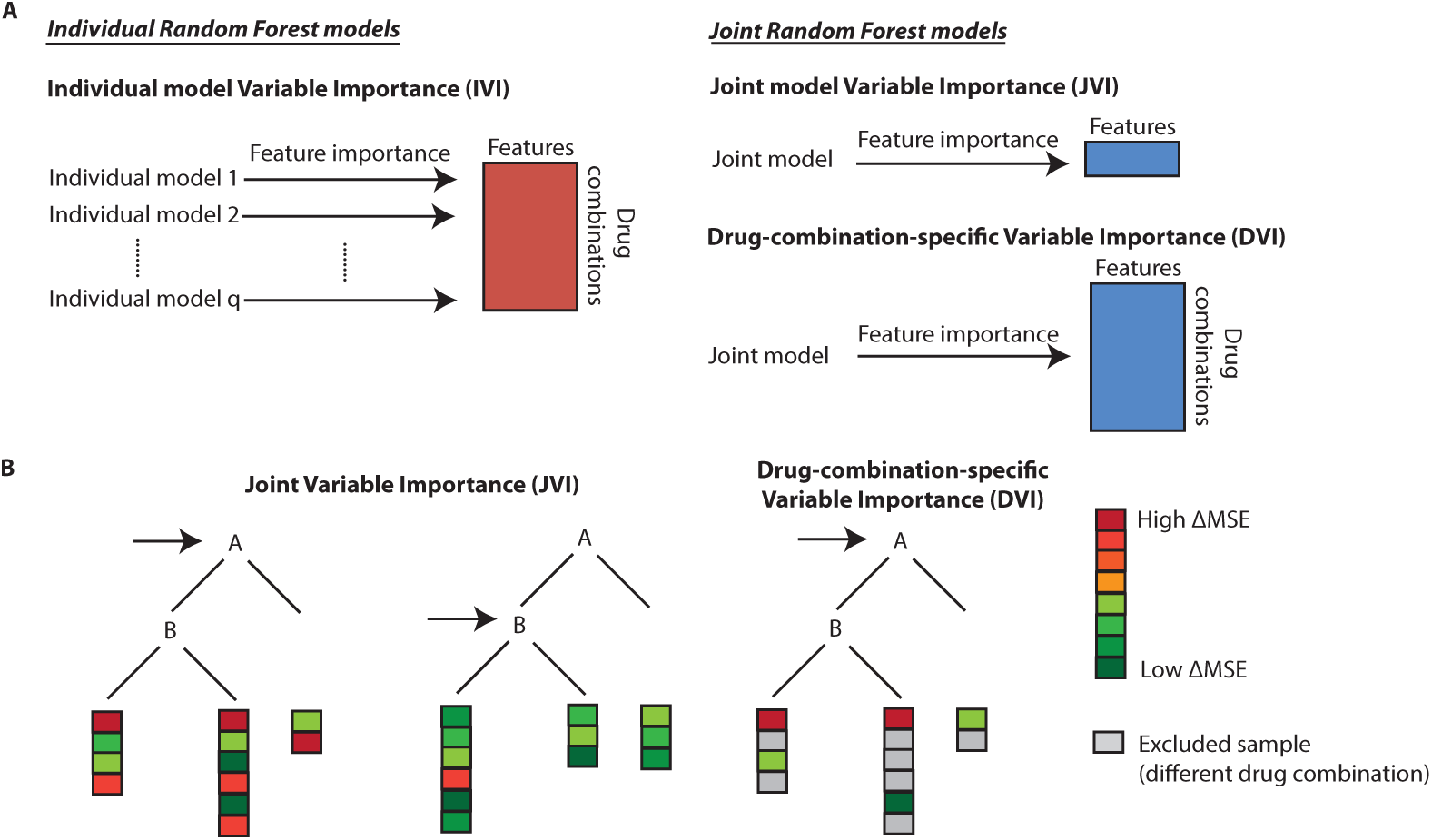
Overview of variable importance scores used in this work. A: Illustration of variable importance scores. Applying the variable importance score normally used with Random Forests to the Joint Random Forest model yields the Joint model Variable Importance (JVI), a measure of variable importance across all drug combinations. Applying the variable importance score normally used with Random Forests to Individual Random Forest models yields the Individual model Variable Importance (IVI), a measure of variable importance per variable and per drug combination. In order to obtain a variable importance for each drug combination using the Joint Random Forest model, we propose the Drug-combination-specific Variable Importance (DVI). B: Illustration of the JVI and the DVI in a single decision tree from the random forest. For both variable importance scores, the importance is assessed by permuting the values of the given variable (a permuted variable is indicated by a horizontal arrow here) and then calculating for each sample (a sample is indicated by a box at the bottom of the tree) the difference between the permuted and unpermuted errors. In the given example, variable A is more important than variable B, as indicated by the higher difference in error (ΔMSE) when permuting variable A.

**Fig 4.**
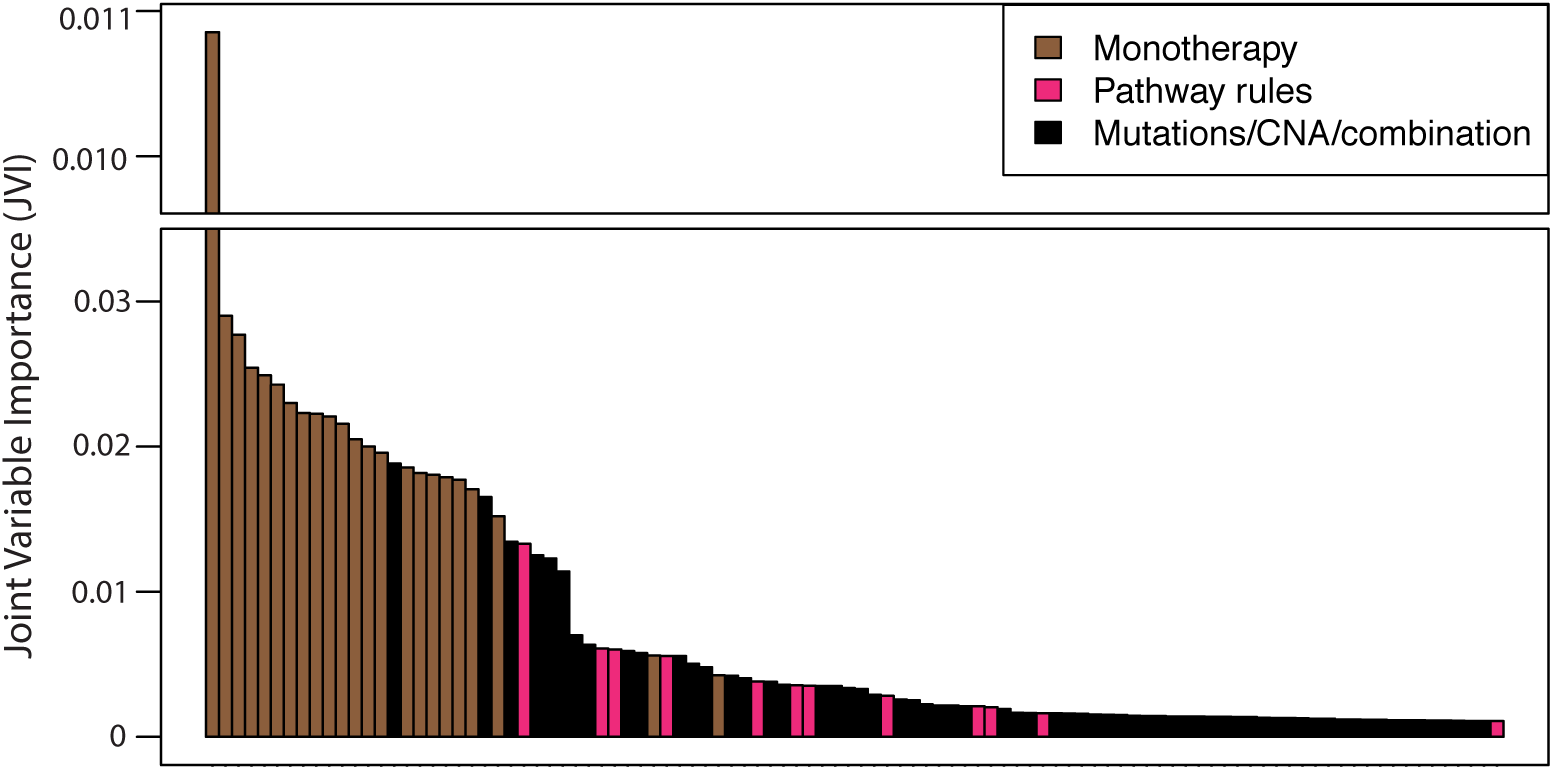
Top 100 highest JVI scores. Joint model Variable Importance (JVI) of the top 100 (out of 664) overall most important variables (for all drug combinations together).

A major drawback of the JVI is that the associations cannot be traced back to specific drug combinations, limiting its use for finding biomarkers of synergy for specific drug combinations. For example, the highest-ranked molecular data variable was mutations in *ATAD5*, which we are unable to link to a specific drug combination using the JVI. To identify biomarkers, we needed a drug-combination-specific variable importance score. Although this could be achieved by computing Random Forest VI scores for the individual models, referred to as Individual model VI score (IVI) (Fig 3A), we preferred to do this using the Joint Random Forest model because of its superior predictive performance. Thus, we needed to define a measure of variable importance per drug combination and per variable using the Joint Random Forest model.

### Drug-combination-specific Variable Importance identifies biomarkers of synergy

To identify biomarkers for a specific drug combination using the Joint Random Forest model, we developed a Drug-combination-specific Variable Importance score (DVI) (Fig 3A). The DVI determines the contribution of each variable to the prediction in the same way as the original Random Forest VI score, but only considers the samples from one drug combination at a time (Fig 3B). To evaluate the DVI, we created a simulated dataset in which we engineered a biomarker with two parameters: 1) *e*: the effect size of the association of the biomarker with synergy; and 2) *d*: the number of drug combinations for which this biomarker was engineered to be associated with synergy. As expected, increasing either one of these enhances the ability of the DVI to identify the biomarker (Fig 5B).

**Fig 5.**
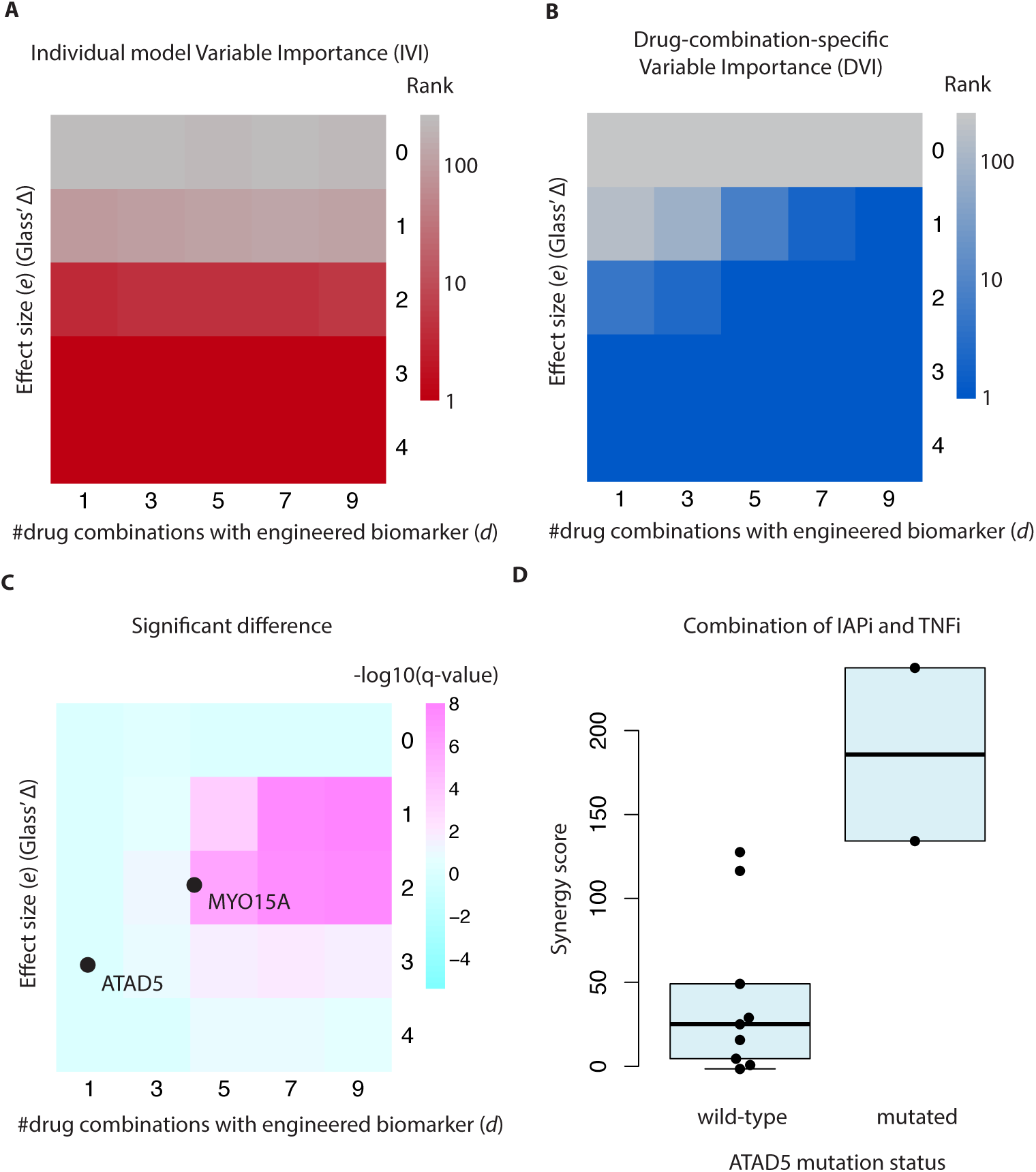
Associations using the IVI and the DVI. Associations using the Individual model Variable Importance (IVI) and the Drug-combination-specific Variable Importance (DVI) A&B: Heatmap showing the median rank of the engineered biomarker in the simulated dataset, stratified by effect size (*e*) and number of drug combinations for which the biomarker was engineered to be associated with synergy (*d*), using either (A) the IVI or (B) the DVI. C: Heatmap showing for which *e* (effect size) and which *d* (number of drug combinations for which the biomarker was engineered to be associated with synergy) the DVI is significantly better (indicated in pink) than the IVI at retrieving the association in a simulated dataset. Examples used in this paper from the DREAM data (associations with *MYO15A* and *ATAD5*) are indicated in this plot based on their effect size and the number of drug combinations in which we observe them. D: Synergy score for a combination of an IAPi and an TNFi, stratified by *ATAD5* mutation status.

We then used the simulated dataset to compare the DVI to the Individual model Variable Importance (IVI, Random Forest VI score on individual models). This showed that the ability of the IVI to identify the engineered biomarker is correlated with the effect size (*e*), but not with the number of drug combinations (*d*) (Figure 5A). This is expected, since the Individual Random Forest models (underlying the IVI scores) do not share information across different drug combinations. Hence, increasing d has no effect on the model’s ability to recover the biomarker. Interestingly, we found that biomarkers with a sufficiently large effect size are identified by both the IVI and the DVI. We found that the DVI is significantly better than the IVI at identifying the biomarker in scenarios where e is small and d is high (Figure 5C).

These findings were reflected in the DREAM data. For example, ranking the associations by their DVI, the highest-ranking molecular data variable was the association of *ATAD5* mutation status with synergy between IAP inhibitors and TNF inhibitors (Figure 5D). As *ATAD5*, IAP and TNF are all part of the apoptosis pathway, this illustrates that the DVI is able to identify interesting associations. Given the large effect size, it is not surprising that this association is ranked high for this drug combination by both the DVI (ranked #3) and the IVI (ranked #1). Using the effect size from this association (Glass’ Δ = 2.8) and assuming that the biomarker is not shared with any other drug combinations, we related this example to the simulated data and found that this example falls in the region where the DVI has little added value (Fig 5C).

In addition, we identified a biomarker (*MYO15A* mutations) that is exclusively identified by the DVI. Given that we observed this association in four related drug combinations and an average Glass’ Δ of 0.87, this example indeed falls in the region where the DVI improves over the IVI (Fig 5C).

### MYO15A mutations associate with synergy between an ALK / IGFR dual inhibitor and PI3K pathway inhibitors in triple-negative breast cancer

We set out to identify biomarkers that were exclusively identified by the DVI. To this end, we decided to focus on the drug combinations containing IGFR inhibitors, for which the joint model obtained the largest increase in predictive performance over the individual models (Fig 2C). Using the DVI to rank all non-monotherapy variables for each of these drug combinations, we found that *MYO15A* mutations had the highest average rank. The association between *MYO15A* mutations and synergy was strongest in combinations of an ALK / IGFR dual inhibitor with PI3K pathway inhibitors (two AKT inhibitors, one PIK3CB / PIK3CD inhibitor and one mTOR inhibitor), hence we decided to further focus on these. These combinations were tested in 23 breast cancer cell lines, of which 20 were triple negative.

The association between *MYO15A* and the synergy score was strongest in the combination containing the PIK3CB / PIK3CD inhibitor (Fig 6A). For the other combinations, the effect was in the same direction and hence in support of this association. Even though these effects would not have been considered significant individually, the model leverages the information across the drug combinations.

**Fig 6.**
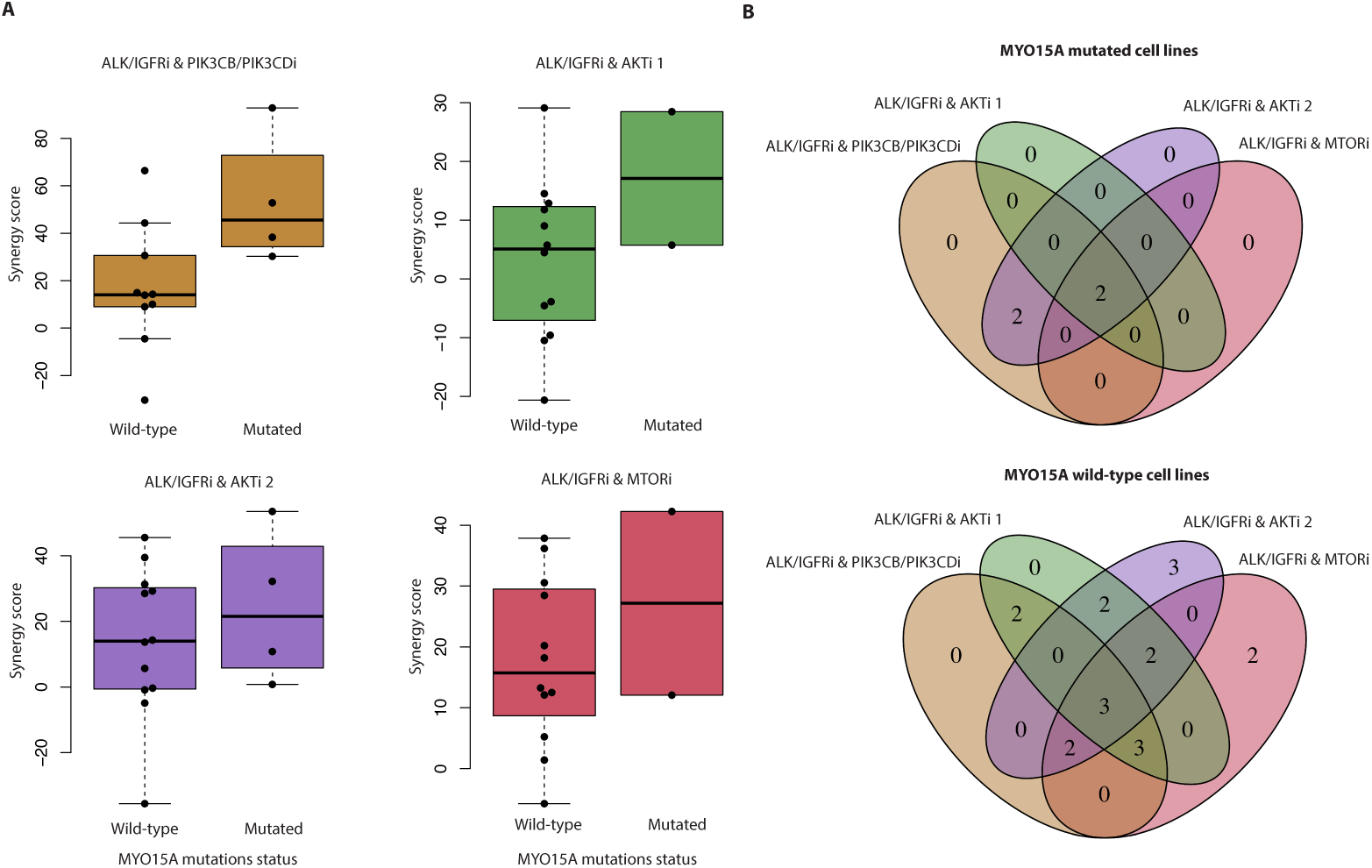
MYO15A mutations associate with synergy between ALK / IGFR dual inhibitors and PI3K pathway inhibitors. A: Synergy scores for combinations of ALK / IGFR dual inhibitors and PI3K pathway inhibitors, stratified by *MYO15A* mutation status. B: Venn diagrams illustrating the overlap in cell lines screened for these four drug combinations. Top panel: *MYO15A* mutant cell lines, bottom panel: *MYO15A* wild-type cell lines.

Of the cell lines in which these combinations have been screened, only five cell lines (two *MYO15A* mutant lines, three wild-type lines) were screened in all combinations; the remaining 18 cell lines (two mutant, 16 wild-type) were not (Fig 6B). Hence, by combining these different combinations, the sample size is effectively increased to 23 cell lines. Altogether, this illustrates how the DVI can be used to identify biomarkers of synergy using the joint Random Forest model.

## Discussion

Drug combinations are of great interest in cancer care, as they can increase treatment efficacy. However, without specific biomarkers, it is difficult to predict which drug combinations will have a synergistic effect in a given patient. Most current approaches for identifying biomarkers of single drug response fit a separate model for each drug. We have shown that such an approach does not obtain good prediction performance for predicting synergy in the DREAM data, likely due to the low sample size. To alleviate this limitation, we used multi-task learning to leverage the information contained in several drug combinations. Compared to previous work [9,13,14,19], our model has the advantage that it is not a ‘black-box method’ and hence can identify biomarkers.

In our models, we found that monotherapy data are important for predicting synergy. Recently, Gayvert *et al.* (2017) [7] have analyzed a similar drug synergy screen, in which they report the same, but do not offer a rationale. We believe that the link between monotherapy and synergy could be attributed to both biological and technical reasons. A biological explanation may be that a small reduction in viability using monotherapy can be evidence of target engagement by the drug, which is required for synergy. On the other hand, the high variable importance of monotherapy can also be technical. When one of the drugs is very potent (e.g. kills 80% of the cells by itself), the expectation is that the combination will kill most cells even if the effect is only additive. Hence, detecting the difference between synergy and additivity would become very difficult in this scenario, as this difference may not exceed the noise level. We note that both scenarios are supported by the data: some drug combinations show positive correlation between monotherapy sensitivity and synergy (corresponding to the first scenario), while others show a negative correlation (supporting the second scenario) (S3 Text).

Another interesting observation is that, on average, using a joint or individual Random Forest model leads to similar predictive performance for drug combinations containing DNA damaging agents. This may reflect previous observations that monotherapy response to DNA damaging agents is notoriously hard to predict [1,18], which might also apply to synergy prediction.

An interesting extension of our method would be the inclusion of variables specific to single drugs or drug combinations, such as chemical structures. These variables could be encoded for each drug or drug combination, similar to the way information is currently encoded specifically for cell lines. While such variables would be correlated with the drug combination indicator variables used in the current model, they could contain additional information, for example when two drug combinations are chemically similar to each other. As chemical information was only available for a subset of the drugs in the DREAM data, we were unable to use this information efficiently. This could be investigated in future work.

In summary, we have presented a method that circumvents the problem of low sample sizes by combining information across drug combinations. In contrast to previous work, our method allows for the identification of biomarkers. With the large number of possible drug combinations, many future drug combination screens are likely to be performed in a small number of cell lines. We believe that our approach can aid to identify biomarkers specifically in such screens.

## Methods

### Individual and joint prediction models

Predictive models are typically trained per drug combination (Fig 1A). We refer to these models as ‘individual models’. In this work, we propose a ‘joint model’, which is simultaneously trained on data from all drug combinations (Fig 1B). Such an approach can be viewed as multi-task learning, where each drug combination represents a task.

The joint model takes an augmented matrix **X**^∗^ as input, in which each sample represents a cell line, drug combination pair. We used an indicator variable to code the different drug combinations. The remaining variables are either:

1. Private to a cell line, drug combination pair (i.e. monotherapy and pathway rules), and hence unique to every sample in **X**^∗^.
2. Private to a cell line only (i.e. mutation and CNA data), and hence repeated across drug combinations.
3. Private to a drug combination only (e.g. chemical structures), and hence repeated across cell lines (though apart from the indicator variables to code the different drug combinations, none were used in this work).

These three categories are visualized in Fig S2 in purple, green and blue respectively.

The response vector *y*^∗^ was defined as the concatenation of the response values, such that each sample corresponds to a cell line, drug combination pair. The resulting input data **X***^∗^* can be fitted onto *y*^∗^ using standard machine learning algorithms. In this work, we have compared three different algorithms:

- Elastic Net [20], as implemented in the R package glmnet [6], with *α* set to 0.5 and *λ* optimized in a nested cross-validation loop.
- SVM [3] with RBF kernels, as implemented in the Python package scikit-learn [16], optimizing the hyper-parameters *C* over [50, 100, 200, 300] and *γ* over [0.001, 0.0001, 0.0005, 0.00005, 0.00001] in a nested cross-validation loop.
- Random Forest [10], as implemented in the Python package scikit-learn, using default parameters.

We compared these joint models to ‘individual models’, which are trained per drug combination and contain the same variables, except the drug combination indicator variables (which are constant within a given drug combination). Predictive performance was assessed using 2-fold cross-validation with the ‘primary score’ (a weighted average of the correlation between the observed and predicted synergy scores) as described in Menden *et al.* (2017) [14] as endpoint. The exact definition of the primary score is

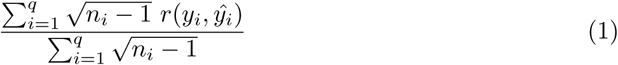

where *q* is the number of drug combinations, *n*_*i*_ the number of cell lines in drug combination *i, r* a function that computes the Pearson correlation, *y*_*i*_ the synergy scores for drug combination *i* and *ŷ*_*i*_ the predicted synergy scores for drug combination *i*. For a fair comparison, all different models (individual and joint; Elastic Net, SVM and Random Forest) were tested using the same cross-validation folds.

### Simulation study to compare the predictive performance of individual and joint Random Forest models

To assess in which situations the joint model outperformed the individual one, we simulated a dataset with 14 cell lines, 497 variables and 10 drug combinations. This dataset closely follows the DREAM dataset in terms of number of samples (median number of cell lines per drug combination is 14) and number of variables (497). For the number of drug combinations, we have limited ourselves to 10, to reduce the computational burden.

For each entry in the input matrix **X**, a value was drawn from a Bernoulli distribution with *p* = 0.5. For each entry in the response vector *y*, a value was drawn from a standard-normal distribution. We then engineered an association between the synergy scores (for all drug combinations in the simulation) and variable *j* by setting *y* = *y* + 2**X***_j_*, resulting in an average effect size of two.

Subsequently, we trained an individual and a joint Random Forest on these data. We evaluated the predictive performance by generating a separate test set, using the same characteristics as the ones used to create the training data. The performance was measured using the ‘primary score’ described above.

To study the effect of sample size on the predictive performance, we varied the sample size between 14, 25, 50 and 100. Likewise, to study the effect of the number of variables, we varied the number of variables between 125, 250, 497 and 1000.

### Variable importance measures

The basic idea of the permuted variable importance [2] is that a variable is considered to be important if it has a positive effect on predictive performance. The importance of a variable **X***_j_* is evaluated by, for a given tree in the forest, calculating the prediction accuracy of the tree on out-of-bag (OOB) samples before and after permuting the values of variable **X***_j_*. The (positive) difference between the two accuracy values is the permuted variable importance for the given tree. The permuted variable importance for the Random Forest is calculated by taking the mean difference in prediction accuracy over all trees in the forest.

In this work, we used the permuted variable importance in two different ways (Figure 3A). When applied to the Joint Random Forest model, we get a global overview of variable importance, which we call the Joint model Variable Importance (JVI). When applied to Individual Random Forest models, we get variable importance scores per variable and per drug combination.

Additionally, to get variable importance scores per drug combination from the Joint Random Forest model we defined the Drug-combination-specific Variable Importance (DVI). This is a modified version of the permuted variable importance, in which the prediction accuracy is calculated using only the samples that correspond to the combination of interest (Figure 2B). For the Random Forest, this accuracy is calculated using a weighted mean square error, in which OOB samples belonging to the combination have weight 1 and all other samples have a weight of 0. A Python implementation of the DVI is available from https://github.com/NKI-CCB/multitask_vi/.

### Simulation study to compare the Individual Variable Importance and the Drug-combination-specific Variable Importance

We generated a simulated dataset as above (14 cell lines, 497 variables and 10 drug combinations; values in **X** drawn from a Bernoulli distribution with *p* = 0.5; and values in *y* drawn from a standard-normal distribution). We then engineered an association between drug combination *i* and variable *j* by setting *y*_*i*_ = *y*_*i*_ + *e***X***_j_*, where *e* is the effect size. To compare the individual and joint models in different configurations, we varied the effect size of the association between *e* = [0, 1, 2, 3, 4] and we varied the number of drug combinations in which the association occurred between *d* = [1, 3, 5, 7, 9], leading to a total of 25 configurations. To estimate the variability in each configuration, we repeated the aforementioned process 50 times for each configuration, leading to a total of 1250 datasets. Each of the 1250 datasets was analyzed using:

- A Joint Random Forest, followed by DVI to rank the variables per drug combination.
- 10 Individual Random Forest models, followed by IVI to rank the variables per drug combination.

For each of the parametrizations (*e* and *d*), we determined:

- The median rank of the engineered association using a Joint Random Forest.
- The median rank of the engineered association using an Individual Random Forest.
- The significance of the difference between these two medians, using a Wilcoxon signed-rank test.

When *d* = 1, we determine median rank of the engineered association using the 50 repeats. When *d* = 3, we use the ranks for the 50 repeats in each of the 3 drug combinations in which the association was engineered, essentially yielding 150 repeats. In general, for each parameterization we determine the median rank of the engineered association using the 50*d* repeats.

We determined the significance of the difference between the individual and the joint model as follows. For each repeat and for each parametrization of *e* and *d*, we determined the median rank of the association across the d drug combinations in which the association was engineered. For each parametrization of *e* and *d*, we then determined the significance using a Wilcoxon signed-rank test (across the 50 repeats). The resulting p-values were corrected for multiple testing using a Benjamini-Hochberg correction.

## Supporting information

**S1 File. List of genes used in copy number alteration analyses.**

**S2 File. List of pathway rules.**

**S1 Text. Data description.**

**S2 Text. Negative correlations in cross-validation results.**

**S3 Text. Association of monotherapy variables with synergy.**

**S4 Text. Description of the pathway rules.**

## Acknowledgments

We would like to thank the AstraZeneca-Sanger DREAM challenge consortium, for organising the DREAM challenge and making the data available. We would like to thank Marlous Hoogstraat for subtyping the breast cancer cell lines. The research leading to these results has received funding from: the European Research Council under the European Union’s Seventh Framework Programme (FP7/2007-2013) / ERC synergy grant agreement n*^◦^* 319661 COMBATCANCER; the Dutch Cancer Society and AACR (SU2C) Dream Team Award (SU2C-AACR-DT1213); and the Netherlands Organization for Scientific Research (NWO, grant Zenith 93512009).

---

**Fig S1.**
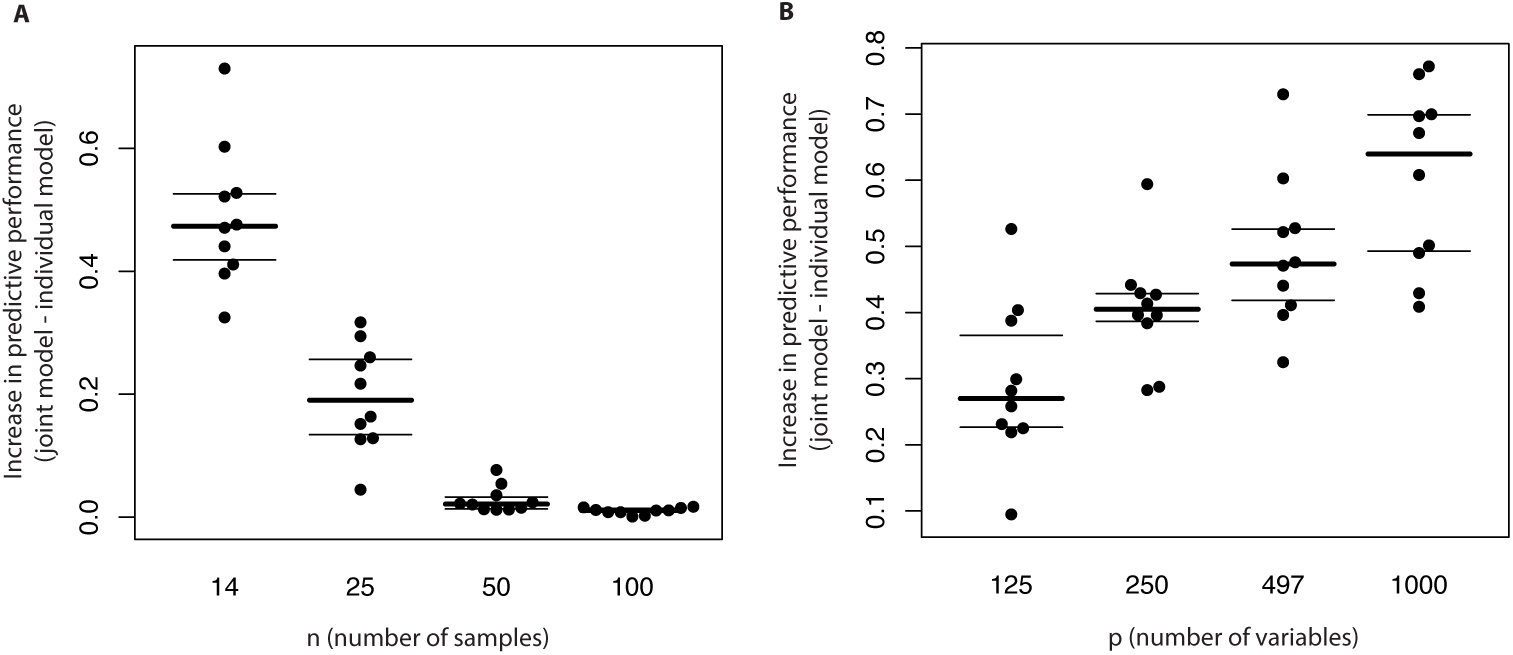
Simulation study to assess under what conditions the joint model outperforms the individual model using Random Forests. A: Difference in predictive performance between the joint and individual model for different values of n (number of samples). B: Difference in predictive performance between the joint and individual model for different values of p (number of variables).

**Fig S2.**
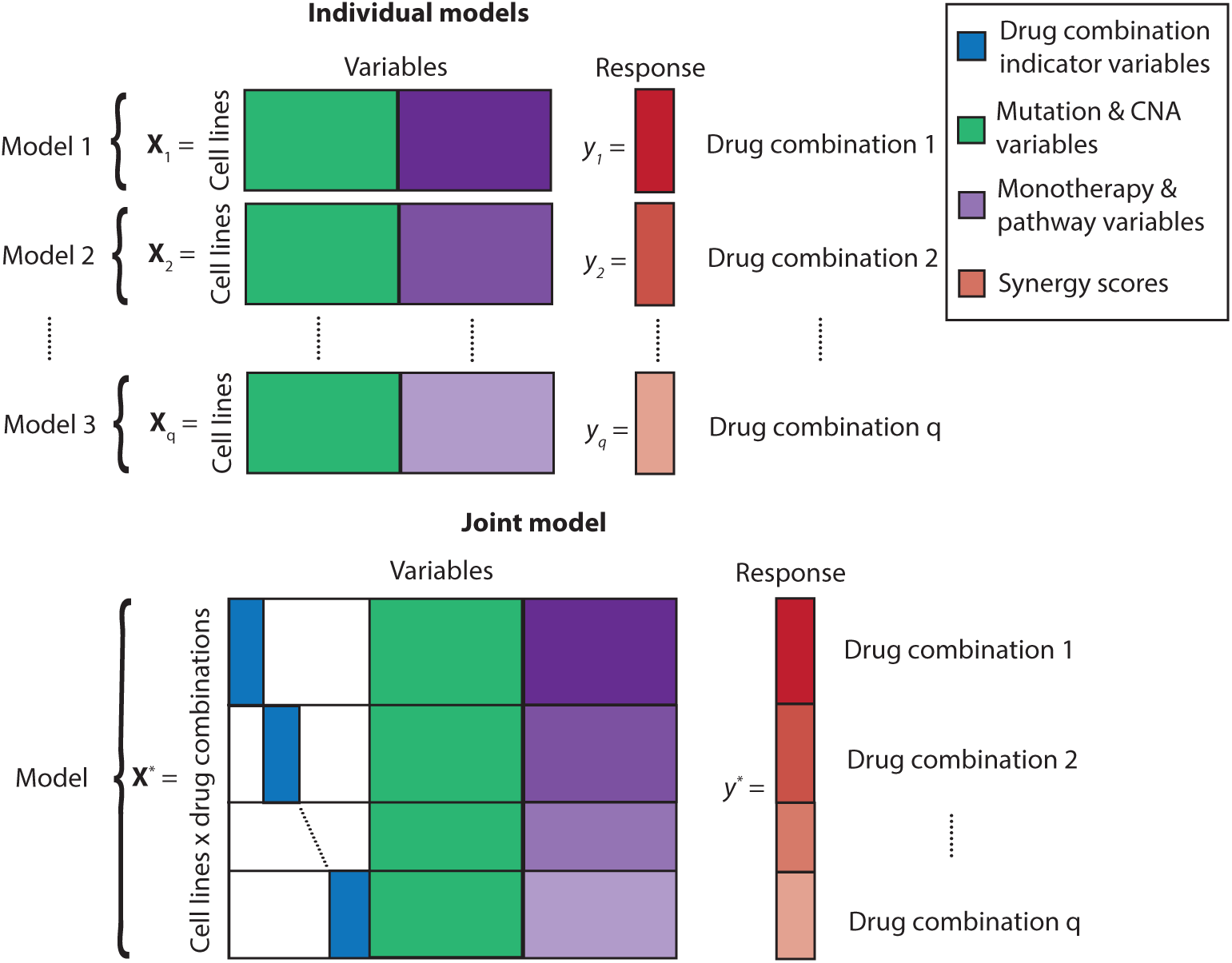
Overview of models used in this work. This figure is similar to Fig 1, but with the added value of showing which variables are private to a cell line, drug combination pair (purple), private to a cell line (green) or private to a drug combination (blue). A: Individual models B: Joint model.

